# Artificial Intelligence Guided Conformational Mining of Intrinsically Disordered Proteins

**DOI:** 10.1101/2021.11.21.469457

**Authors:** Aayush Gupta, Souvik Dey, Huan-Xiang Zhou

## Abstract

Artificial intelligence recently achieved the breakthrough of predicting the three-dimensional structures of proteins. The next frontier is presented by intrinsically disordered proteins (IDPs), which, representing 30% to 50% of proteomes, readily access vast conformational space. Molecular dynamics (MD) simulations are promising in sampling IDP conformations, but only at extremely high computational cost. Here, we developed generative autoencoders that learn from short MD simulations and generate full conformational ensembles. An encoder represents IDP conformations as vectors in a reduced-dimensional latent space. The mean vector and covariance matrix of the training dataset are calculated to define a multivariate Gaussian distribution, from which vectors are sampled and fed to a decoder to generate new conformations. The ensembles of generated conformations cover those sampled by long MD simulations and are validated by small-angle X-ray scattering profile and NMR chemical shifts. This work illustrates the vast potential of artificial intelligence in conformational mining of IDPs.

## Introduction

Artificial intelligence (AI) is gradually overshadowing traditional physics-based approaches^1, 2^, achieving breakthroughs in solving some of the most challenging problems in chemistry and physics. For example, a deep neural network has obtained nearly exact solutions of the electronic Schrodinger equation for small moleucle^3^. Another recent breakthrough is the prediction of three-dimensional structures of proteins by neural network-based methods, Alphafold^4^ and RoseTTafold^5^. With problems facing structured proteins being solved by these and other AI-based methods^6–9^, a new frontier is presented by intrinsically disordered proteins (IDPs). Instead adopting well-defined three-dimensional structures, IDPs readily access vast conformational space. Here we report on the development of a generative AI model to mine the conformational space of IDPs.

IDPs, accounting for 30% to 50% of proteomes, perform many essential cellular functions including signaling and regulation, and are implicated in numerous human diseases^10, 11^. In particular, polyglutamine expansion is associated with Huntingtin’s and other diseases^12^. Amyloid-beta peptides, including Aβ40, are linked to Alzheimer’s disease^13^. The cell division machinery of *Mycobacterium tuberculosis*, the causative agent of tuberculosis, contains a number of membrane proteins, including ChiZ, with disordered cytoplasmic regions^14, 15^. The functional and disease mechanisms of these and other IDPs remain unclear, in large part because we lack knowledge of their conformational ensembles in various states (e.g., in isolation, in aggregation, and bound with interaction partners).

The vastness of IDPs’ conformational space poses significant challenges. Experimental techniques are limited to probing some aspects of the conformational space. For example, small-angle x-ray scattering (SAXS) provides information on the overall shapes and sizes of IDPs^16^, whereas NMR properties, such as secondary chemical shifts, carry residue-specific information but still vastly under-represent the degrees of freedom of IDPs^17^. Molecular dynamics (MD) simulations offer an attractive approach for IDPs, with an atomic representation for each conformation, but the simulation time that can be presently achieved, which directly determines the extent of conformation sampling, is largely limited to 10s of μs. The conformational ensembles of the 64-residue cytoplasmic disordered region of ChiZ (referred to simply as ChiZ hereafter) sampled by multiple replicate simulations, totaling 36 μs in solution and 38 μs at membrane, have been validated by SAXS and NMR data^14, 15^. While we cannot answer whether 10s of μs of simulations are really long enough, we do know that shorter simulations are insufficient. For example, Kukharenko *et al*.^18^ have shown that the conformations of a 22-residue fragment of α-synuclein sampled in 1 μs represent only a small subset of the ensemble collected from 13 μs of “expansion” simulations. The latter are a large number (200) of short simulations (30-100 ns) started from sparsely populated regions in a two-dimensional embedded space (via sketch-map embedding). How to exhaustively cover the conformational space of IDPs without an inhibitory amount of computational time remains an open question.

For structured proteins, autoencoders have been developed to represent structures in two-dimensional latent spaces and reconstruct the structures back in Cartesian coordinates^6, 8^. In another recent study^9^, an autoencoder was trained to project the inter-residue distances of the ribose-binding protein into a two-dimensional latent space. The open and closed states of the protein were found to occupy separate regions in the latent space. The authors linearly interpolated points from these two states and decoded the interpolated points into inter-residue distances that represent conformations on the transition paths between the open and closed states. The inter-residue distances from interpolation were finally coupled to an all-atom model to enhance the latter’s conformational sampling in MD simulations. Noé et al.^7^ built Boltzmann generators, which use neural networks to represent protein structures sampled from short MD simulations as a Gaussian distribution in the latent space; points sampled from the Gaussian are transformed back as structures in Cartesian coordinates. In toy problems, the authors demonstrated that points located in different energy wells in conformational space are repacked into a dense distribution with a single peak in the latent space. These and other AI-based methods might potentially be adapted to study IDPs^19^.

Here we present generative autoencoders designed to mine the conformational space of IDPs. The performance of these autoencoders rivals that of expensive MD simulations and is validated by SAXS and chemical shift data. Our work opens the door to modeling IDPs in various functional states.

## Results

We built autoencoders first to represent IDP conformations as vectors in a reduced-dimensional latent space (Fig. 1A). Training of the autoencoders involved reconstructing the conformations from the latent vectors and minimizing deviations from the original conformations. The training datasets consisted of conformations sampled from short MD simulations. We then modeled the latent vectors of the training datasets as multivariate Gaussian distributions (Fig. 1B). By sampling from these distributions for reconstruction, we generated the full conformational ensembles of IDPs (Fig. 1C). These generative autoencoders were built for three IDPs: polyglutamine Q15, Aβ40, and ChiZ, and were validated by their ability to cover all conformations sampled in long MD simulations and to reproduce experimentally measured properties. These IDPs contain 17 (including two capping groups), 40, and 64 residues (denoted by *N*_res_).

**Figure 1.**
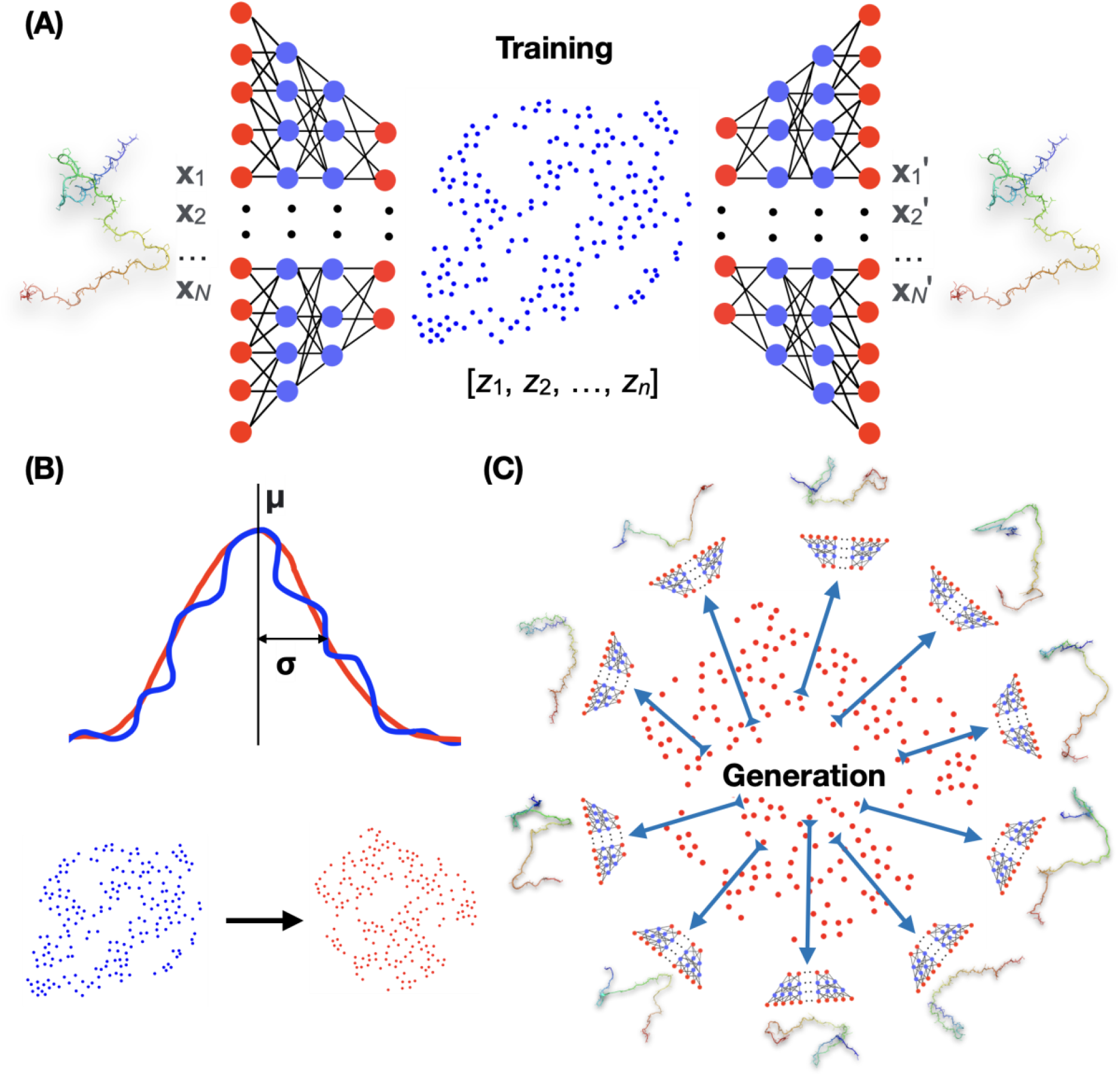
Design of generative autoencoders. (A) Illustration of the architecture of an autoencoder. The encoder part of the autoencoder represents the conformations of an IDP, specified by the Cartesian coordinates [**x**_1_, **x**_2_, …, **x**_N_] of *N* heavy atoms, as *n*-dimensional vectors [*z*_1_, *z*_2_, …, *z*_n_] in the latent space. The decoder then reconstructs the latent vectors back to conformations in Cartesian coordinates, [**x**_1_′, **x**_2_′, …, **x**_N_′] During training, the weights of the neural networks are tuned to minimize the deviation of the reconstructed conformations from the original ones. (B) Modeling of the distribution of the latent vectors (blue) of the training set by a multivariate Gaussian (red). The mean vector and covariance matrix of the Gaussian are those of the training latent vectors. The curves illustrate a Gaussian fit to the distribution of the training data; the scatter plots show a comparison of the training data and the Gaussian model. (C) Generation of new conformations. Vectors sampled from the multivariate Gaussian are fed to the encoder to generate new conformations.

### Representation in a reduced-dimensional space

As a steppingstone to generating new conformations, we first reduced the dimensionality of the conformational space. The original conformations of the IDPs were specified by the Cartesian coordinates of heavy atoms (with truncation for some side chains). The dimension of the conformational space was thus 3*N*, where *N*, denoting the number of heavy atoms included, was 140, 230, and 385, respectively, for Q15, Aβ40, and ChiZ. We chose the dimension (*n*) of the latent space for each IDP to be 0.75*N*_res_, or 13 for Q15, 30 for Aβ40, and 48 for ChiZ.

Conformations for training and testing the autoencoders came from multiple μs-long MD simulations^14, 20^. We collected 95,000, 140,000, and 145,000 frames, respectively, at 10-ps intervals for Q15 and 20-ps intervals for Aβ40 and ChiZ from each replicate run; the numbers of replicate runs were 2, 4, and 12, respectively. An initial portion (e.g., 10%) of each run was taken as a training set whereas the remaining portion was the test set. The accuracy of an autoencoder was assessed by the root-mean-square deviation (RMSD) between each test conformation and its reconstruction. These RMSDs were averaged for the entire 100-fold diluted test set (comprising frames saved at 1-ns intervals for Q15 and 2-ns for Aβ40 and ChiZ). As adjacent frames in MD simulations tend to have similar three-dimensional structures, the dilution serves to reduce redundancy of the test set. The reconstruction RMSD results are shown in Fig. 2.

**Figure 2.**
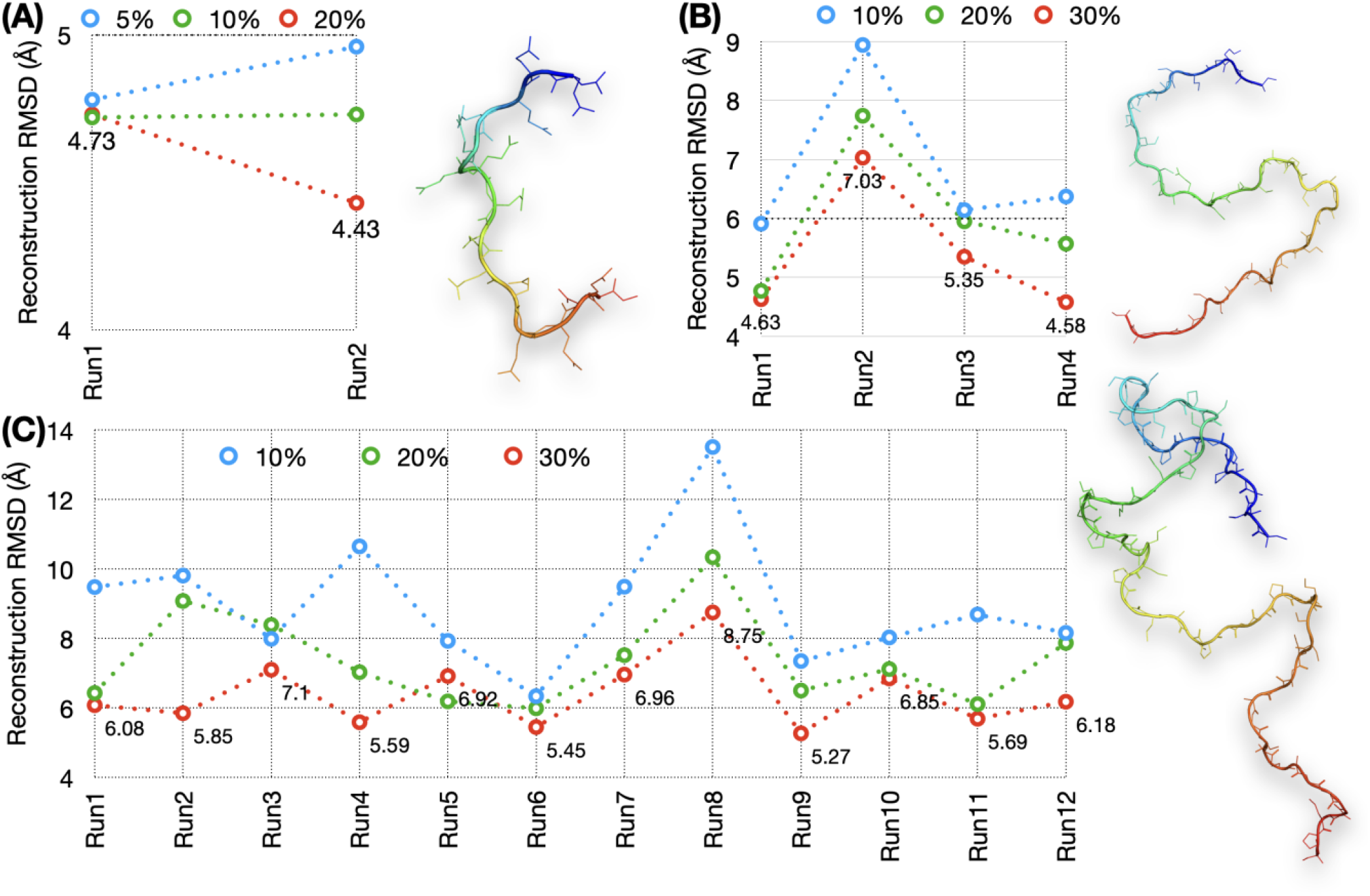
Reconstruction results. Average reconstruction RMSDs at different sizes of the training sets sampled from replicate MD runs. (A) Q15 at 5%, 10%, and 20% training sizes from two runs. (B) Aβ40 at 10%, 20%, and 30% training sizes from four runs. (C) ChiZ at 10%, 20%, and 30% training sizes from 12 runs. A structure for each IDP is shown.

For Q15, the average reconstruction RMSDs are below 5 Å even when only 5% of the MD simulations (corresponding to 95 ns of simulation time) is used for training (Fig. 2A). When the training size is increased to 10% and 20%, the RMSDs stay around 4.75 Å for run1 but decrease successively from 4.96 Å to 4.73 Å and 4.43 Å. This decrease in reconstruction RMSD with increasing training size is likely because run2 was started from an all α-helical conformation, which mostly melted away over time (Fig. S1A). For Q15, we chose autoencoders trained at the 10% size for generating new conformations.

For Aβ40, training with the first 10% of the MD simulations results in reconstruction RMSDs of 6.4 ± 1.3 Å (mean ± standard deviation among four MD runs) (Fig. 2B). The reconstruction RMSDs decrease to 6.0 ± 1.4 Å with a 20% training size and further to 5.4 ± 1.1 Å with a 30% training size. The higher RMSD of run2 is probably due to more compact initial conformations (Fig. S1B). For this IDP we chose a 20% training size for generating new conformations.

Reconstruction becomes more challenging as the IDP size increases. This is already apparent when Aβ40 is compared to Q15, and is much more so for ChiZ, where training with 10% of the MD simulations results in reconstruction RMSDs at 8.3 ± 1.1 Å for 10 of the 12 MD runs, and > 10 Å for the other two runs (Fig. 2C). Still, the reconstruction RMSDs decrease to 7.4 ± 1.3 Å with a 20% training size and further down to 6.4 ± 1.0 Å with a 30% training size. For this larger IDP, we chose 30% training size (corresponding to 870 ns of simulation time) for generating new conformations.

To check whether the dimensions of the latent space chosen according to 0.75*N*_res_ were adequate, we trained autoencoders with a 200-dimentional latent space. The reconstruction RMSDs improve for Q15 and Aβ40, but not for ChiZ (Fig. S2). So increasing the latent-space dimension does not necessarily improve accuracy, especially for the larger, more challenging IDPs, in reconstruction (or in generating new conformations; see below).

### Multivariate Gaussian models in latent space

The conformational ensembles of IDPs are broad and difficult to model^14^. A possible crucial benefit of representing the conformations in the latent space is that, due to the reduced dimensionality, the distribution of the latent vectors would be more compact and therefore easier to model. To assess this expectation, we calculated histograms in two-dimensional subspaces of the latent space. For each autoencoder, about one half of the encoder output values were consistently at or near zero, thereby further reducing the effective dimension of the latent space. We only calculated histograms for pairs of nonzero output neurons.

For the run1 training set of Q15, only 7 of the 13 output neurons were nonzero, resulting in 21 possible pairs. In Fig. S3, we display the histograms of 10 pairs calculated for the training (10% size) and test datasets. These histograms are indeed compact. Moreover, the counterparts of the training and test sets look very similar, with only minor differences for one or two pairs. For example, in the (9, 11) pair, the histogram of the training set is somewhat broader than the counterpart of the test set. The significant overlap between the distributions of the training and test sets in the latent space explains the good performance of the autoencoder in reconstruction.

The autoencoder for Aβ40 (run1; 20% training size) had only 15 nonzero output neurons (out of 30). Fig. 3 displays the histograms of 8 nonzero pairs. All of these are single-peaked, and the peak positions are the same for the training and test counterparts in most cases, but with some shift for the (0, 27) pair. The high-level of overlap between the training and test sets allows for the satisfactory reconstruction of Aβ40 conformations reported above. In comparison, for the larger ChiZ, the histograms representing conformations sampled from a single MD run (run1) become irregular in shape (e.g., the (38, 39) pair) and the divergence between the training and test sets becomes prominent (e.g., the (15, 16) and (44, 47) pairs) (Fig. S4). These features exhibited by the distributions in the latent space illustrate the growing difficulty in reconstructing the conformations of larger IDPs.

**Figure 3.**
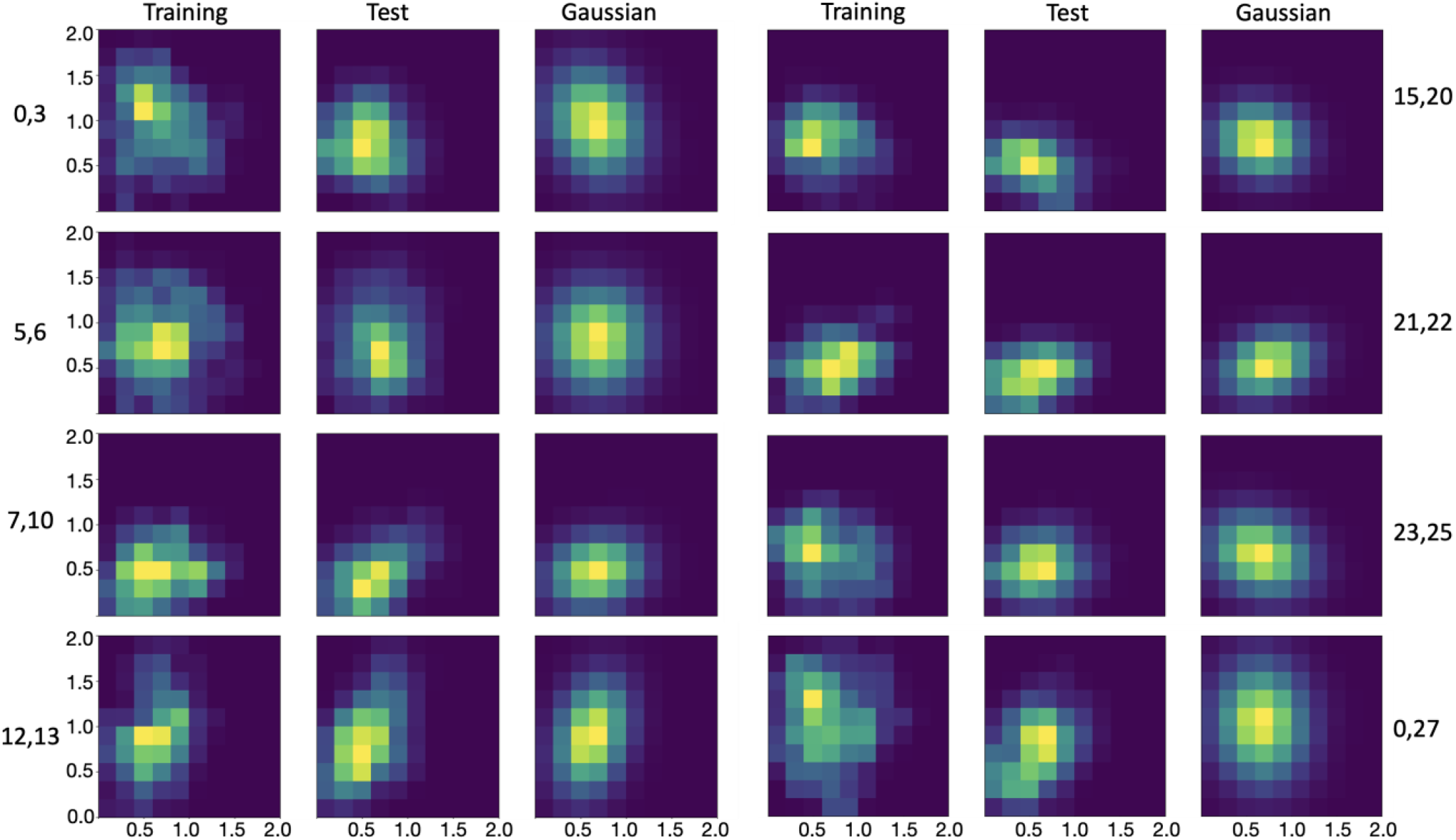
Histograms of Aβ40 in the latent space, calculated from training data, test data, and Multivariate Gaussian. Histograms for pairs of encoder nonzero outputs from run1 are shown as heat maps, with yellow representing pixels with the highest counts and dark blue representing pixels with 0 count.

The compact distributions of Q15 and Aβ40 in the latent space motivated us to model them as multivariate Gaussians. As shown in Figs. S3 and 3, the distributions of the training sets and their multivariate Gaussian models look very similar. More importantly, the multivariate Gaussian models also overlap well with the distributions of the test sets. Indeed, the overlap between the test sets and the Gaussian models is greater than that between the test sets and the corresponding training sets, as illustrated by the (9, 11) pair of Q15 and the (0, 27) pair of Aβ40. Therefore the multivariate Gaussian models seem promising for generating new conformations that are similar to those in the test sets of Q15 and Aβ40. For ChiZ, multivariate Gaussians are inadequate to model the irregular shapes of the single-run distributions in the latent space (Fig. S4).

### Autoencoder-generated conformations of Q15 and Aβ40

By sampling from a multivariate Gaussian in the latent space and using the decoder to reconstructing conformations, we turned the autoencoder into a generative model. The multivariate Gaussian was parameterized on the same dataset for training the autoencoder. For Q15, the training size was 9,500 and the test size was 85,500. The size of the generated set was measured as multiples of the test size (1× = 85,500). For each conformation in the 100-fold diluted test set, we found its best match (i.e., lowest RMSD) in the generated set. We then used the average of the best-match RMSDs for the diluted test set as the measure for the accuracy of the generated set. With the generated sets at size 1×, the average best-match RMSDs of the test sets are 3.59 and 3.58 Å for MD run1 and run2, respectively. As illustrated in the inset of Fig. 4A, a test conformation and its generated best match at 3.88 Å RMSD show very similar backbone traces. Since generating new conformations by the autoencoder is extremely fast, the generated set can be easily expanded. With expanding sizes of the generated set, the average best-match RMSDs show small but systematic decreases, to 3.55 Å at 2×, 3.52 Å at 3×, and 3.51 Å at 4× for run1 (Fig. 4A). The improvement in RMSD occurs because the expanded size of the generated set yields better matches for the test conformations. Conversely, the average best-match RMSDs increase to 3.64 Å when the size of the generated set is reduced to 0.5× and further to 3.79 Å when the generated set is reduced to the same size as the training set (at 0.11×). Increasing the dimension of the latent space to 200 did not improve the accuracy of the autoencoder in generating new conformations (Fig. S5).

**Figure 4.**
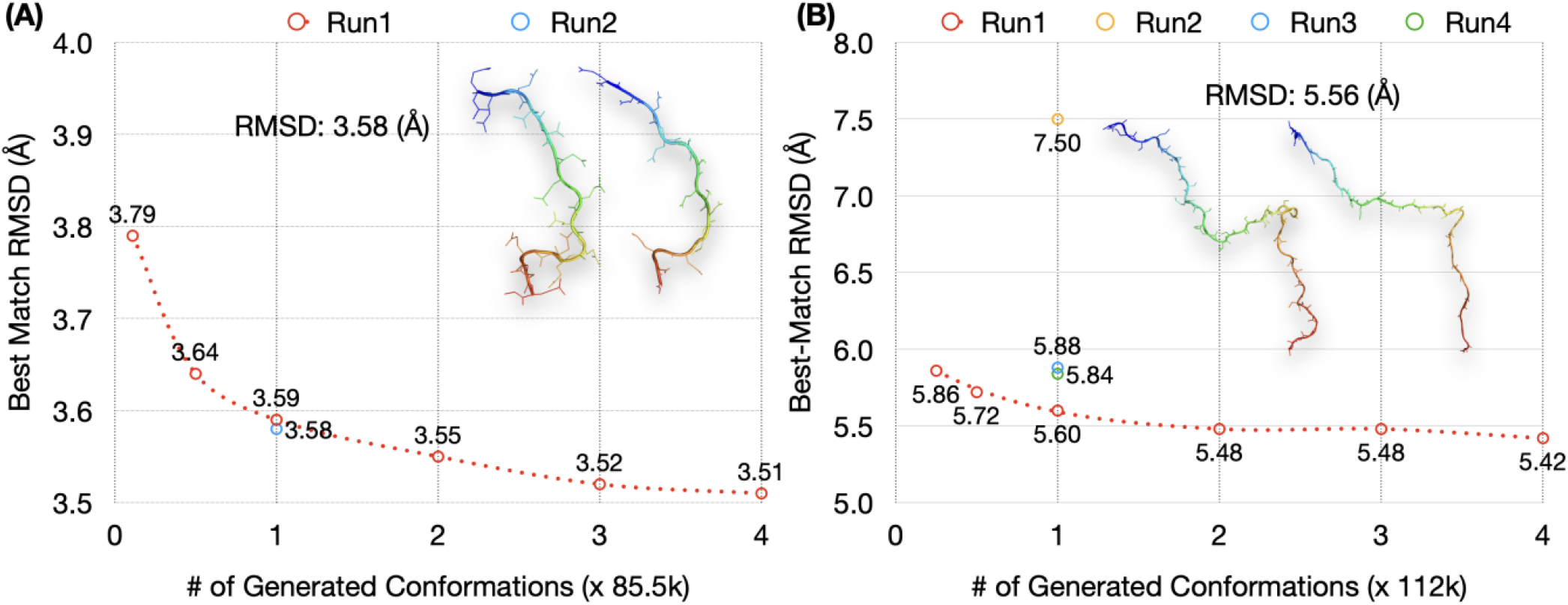
Results for autoencoder-generated conformations of Q15 and Aβ40. The average best-match RMSDs of 100-fold diluted test sets of (A) Q15 and (B) Aβ40, against generated sets at different sizes. The latter sizes are measured in multiples of the test size of each IDP (= 85,500 for Q15 and 112,000 for Aβ40). For run1, results are shown at sizes of the generated set ranging from the training size to 4×. For other MD runs, results are shown at 1×. In the inset of each panel, an IDP conformation and its generated best match, with an RMSD close to the average values at 1×, is compared.

High accuracy is also achieved for generated conformations of Aβ40 on autoencoders trained with 20% (= 28,000 conformations) of MD simulations (Fig. 4B). With the size of the generated sets at 1× (=112,000 conformations), the average best-match RMSDs of the 100-fold diluted test sets are 5.60 Å, 7.50 Å, 5.88 Å, and 5.84 Å, respectively, for MD run1 to run4. A test conformation and its generated best match at 5.56 Å RMSD show very similar backbone traces (Fig. 4B, inset). The higher average RMSD of the autoencoder for run2 in generating new conformations mirrors the poorer performance of this autoencoder in reconstruction (Fig. 2B), and can also be attributed to the overly compact conformations in the training set of this MD run (Fig. S1B). With an expansion of the generated set, the average best-match RMSD shows a slight decrease, to 5.42 Å at 4× for run1 (Fig. 4B). Conversely, the average best-match RMSD increases to 5.72 Å at 0.5× and to 5.86 Å at 0.25× (= size of the training set).

### Autoencoder-generated conformations of ChiZ

We first used a similar protocol to train and test an autoencoder for ChiZ on a single MD run (run1). The training size was 30% or 43,500 and the test size was 101,500. With the generated set at size 1× (= 101,500 conformations), the average best-match RMSD of the 100-fold diluted test set is 7.95 Å (Fig. S6A). Again the RMSD decreases slightly with expanding sizes of the generated set, but is still 7.35 Å even at size 12× (= 1.2 million conformations). The high RMSD of the autoencoder trained on a single MD run is presaged by the inadequate modeling of the training data by a multivariate Gaussian in the latent space (Fig. S4). It is possible that a single MD run may mine a limited region in conformational space, but the regions mined by different MD runs may partially overlap and the combined mining may generate an ensemble that is densely distributed in the latent space. Indeed, when we combine the conformations from 12 MD runs for ChiZ, the histograms in the latent space for both the training set and the test set become compact and have a single peak for all but one (i.e., (9, 14)) of the nonzero pairs (Fig. 5A). The distributions of the training and test latent vectors overlap very well and are also modeled well by the multivariate Gaussian parameterized on the combined training set.

**Figure 5.**
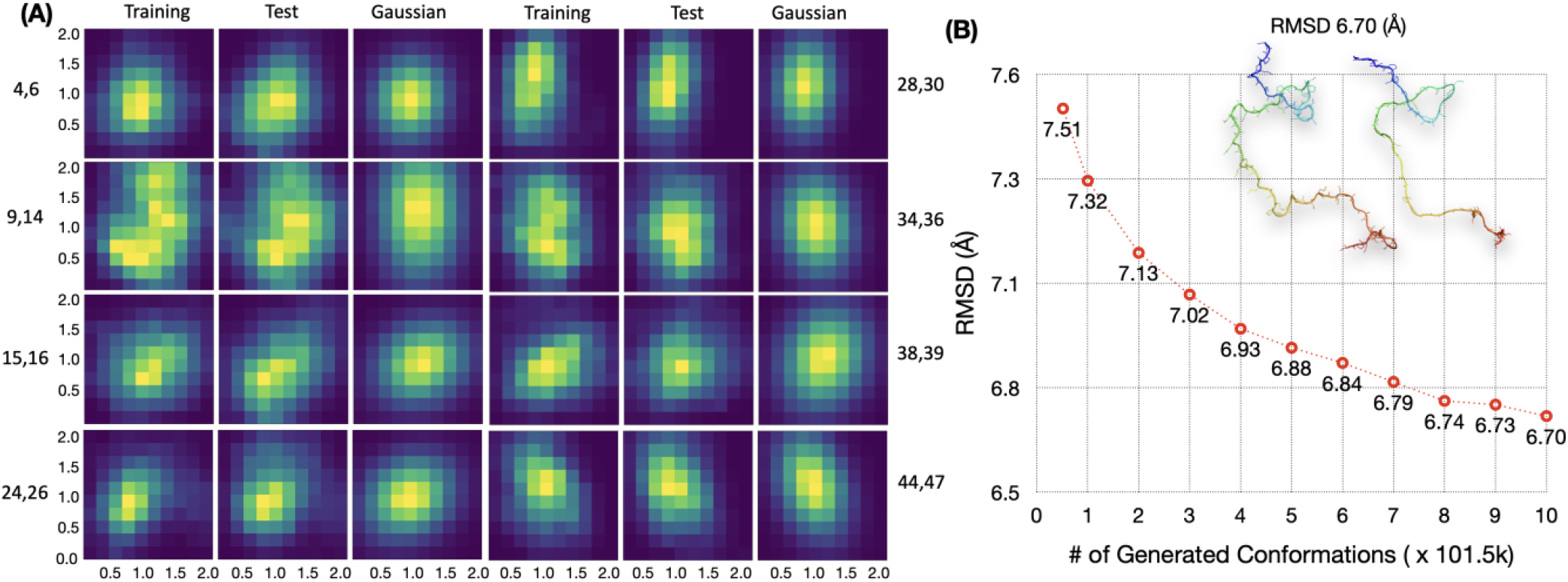
Increased data overlap and prediction accuracy by combining MD runs of ChiZ. (A) Histograms in the latent space, shown as heat maps, with yellow representing pixels with the highest counts and dark blue representing pixels with 0 count. Histograms were calculated for pairs of nonzero elements, using 52,200, 121,800, and 101500 vectors from the training and test sets and the multivariate Gaussian, respectively. The training and test sets were from combining conformations sampled in 12 MD runs; the multivariate Gaussian was parameterized on the combined training set. (B) The average best-match RMSDs of the 1000-fold diluted, combined test set against generated sets at different sizes. The autoencoder was trained on a 10-folded diluted, combined training set (size = 52,200) from all the 12 MD runs. The sizes of the generated sets are measured in multiples of the test size in a single MD run (= 101,500), and range from 0.51× (= training size) to 10×. The inset displays an IDP conformation and its generated best match, with an RMSD of the average value at 10×.

The increase in overlap by combining data from multiple MD runs pointed a way to improve autoencoders. As an initial test, we pooled the generated conformations (each at size 1×) from the autoencoders of the individual MD runs. When compared with this pooled generated set (total size at 12×), the average best-match RMSD of the run1 test set is 7.04 Å (Fig. S6B), which is lower by 0.31 Å than the corresponding value when the generated set is at the same 12× size but produced solely by the run1 autoencoder (Fig. S6A). To take full advantage of the multiple MD runs of ChiZ, we used the autoencoder trained on the combined training set (a total of 52,200 conformations after a 10-fold dilution) to generate new conformations. The generated set at size 1× now gives a best-match RMSD of 7.32 Å for the combined test set (121,800 after a further 10-fold dilution). When the generated set is expanded to a size 10×, the best-match RMSD reduces to 6.70 Å (Fig. 5B). The inset illustrates a pair of conformations, one from the test set and one from the generated set, at this RMSD.

### Further assessment of generated conformations

To properly benchmark the autoencoder-generated conformations, we examined the diversity of the test sets and the similarity between the training and test sets (Table S1). We calculated the RMSDs of each conformation with all others in a diluted test set. The average pairwise RMSDs are quite high even within a single MD run (run1), 6.98 Å for Q15, 11.61 Å for Aβ40, and 18.21 Å for ChiZ, showing that the conformations in each test set are very diverse. As expected, the average pairwise RMSD increases further, to 19.23 Å, for the combined and further diluted test set of ChiZ. The diversity of the test conformations again illustrates the challenge in generating conformations that are close to them.

The neighboring conformations in any MD run have relatively low RMSDs, leading to small best-match RMSDs between conformations in the test sets from single MD runs. The average best-match RMSDs in run1 are 3.71 Å for Q15, 3.83 Å for Aβ40, and 4.83 Å for ChiZ. However, for the combined and further diluted test set of ChiZ, the average best-match RMSD increases to 8.62 Å. The latter value may be viewed as a benchmark for generated conformations to be claimed as neighbors of test conformations. Because the average best-match RMSD for the combined test set against the generated set (at size 10×) is 6.70 Å, or nearly 2 Å below the benchmark, we can claim that all the test conformations in the combined test set have neighbors in the generated set. In other words, the generated set covers the combined test set.

Another benchmark is given by the average best-match RMSD between a test set and the corresponding training set. For run1, values of this benchmark are 3.96 Å for Q15, 6.76 Å for Aβ40, and 10.17 Å for ChiZ. When the comparison is against the generated sets at the sizes of the training sets (shown as the first point in Figs. 4A, 4B, and S6A), the average best-match RMSDs are 3.79, 5.86, and 8.16 Å, respectively, each of which is lower than the counterpart when the comparison is against the training set itself. That is, relative to the training sets, the generated sets provide better matches for the test sets. For Q15 and Aβ40, this outcome is to be expected because of the above observation that the test sets overlap better with the Gaussian models than with the training sets (Figs. S3 and 3). For ChiZ, the combined test set from the 12 MD runs has a best-match RMSD of 8.47 Å against the combined training set, which is 1.7 Å lower than the counterpart for the comparison within run1. This decrease in best-match RMSD confirms the aforementioned increase in data overlap when multiple MD runs are combined (Figs. S4 and 5A). Moreover, the best-match RMSD of the combined test set further reduces to 7.51 Å when the generated set is of the same size as and parameterized on the combined training set (first point in Fig. 5B).

We also inspected more closely the generated conformations that best match test conformations (Fig. 6). As already alluded to, test conformations and their generated best matches show overall similarities in shape and size. However, the generated conformations have significant bond length and bond angle violations and are overly smooth locally, especially for the larger IDPs. Many neural network models are prone to over-smoothing.

**Figure 6.**
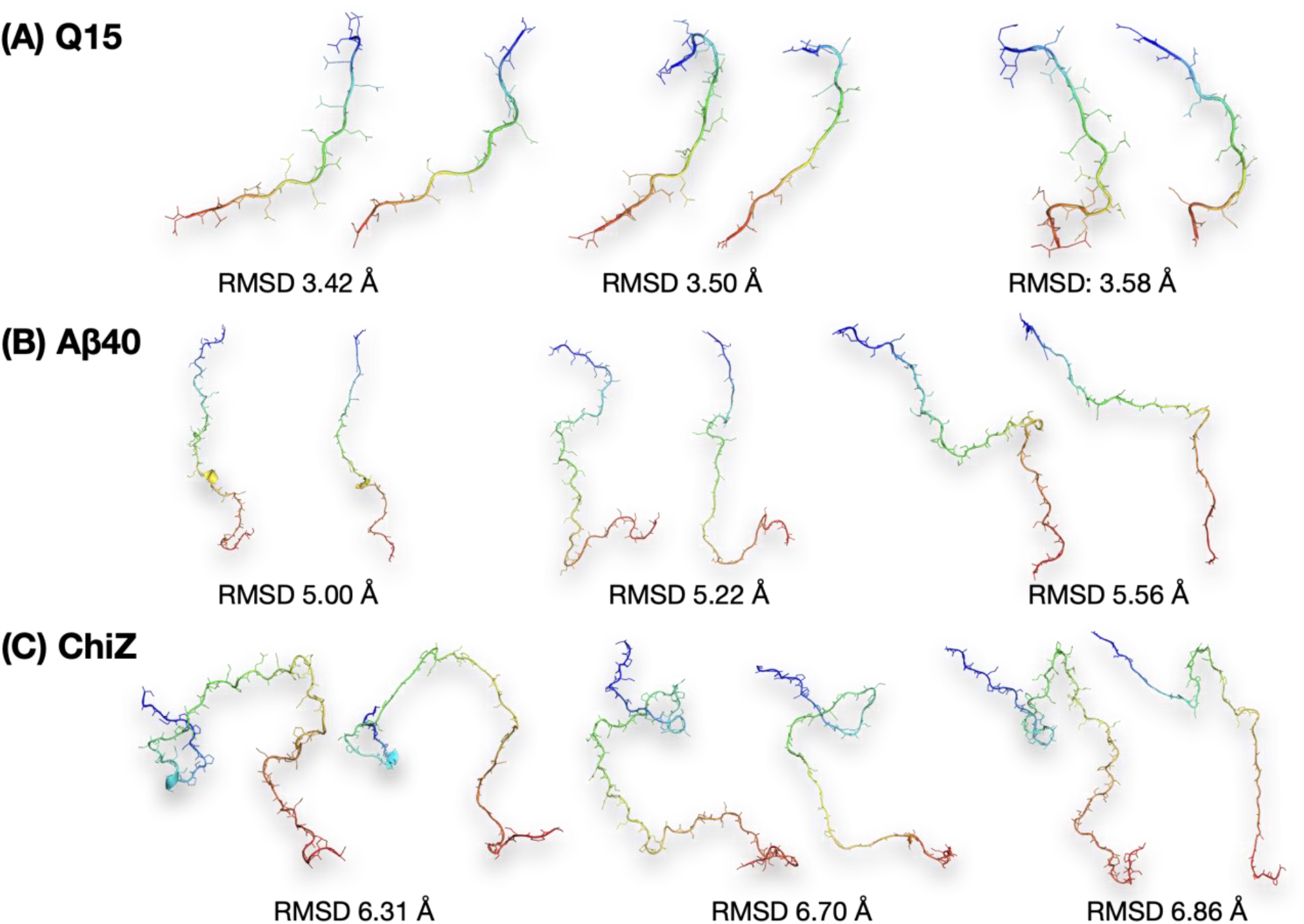
Comparison of test conformations and their generated best matches. (A) Q15. (B) Aβ40. (C) ChiZ. Each IDP is represented by three pairs of conformations with RMSDs around the average best-match value of the diluted test set against the final generated set. In each pair, the left conformation is from the test set and the right conformation is from the generated set. Each chain is colored from blue at the N-terminus to red at the C-terminus.

### Experimental validation of autoencoder-generated ChiZ conformational ensemble

To objectively assess the quality of the autoencoder-generated conformational ensemble, we calculated from it properties that can be measured experimentally. These include SAXS profile and NMR chemical shifts. In Fig. 7, we compare the experimental data for ChiZ^14^ with results calculated from 12,180 conformations collected from the combined test set of the 12 MD runs, and with results calculated from 12,180 conformations generated by the autoencoder trained on the combined training set. As reported previously^14^, the MD simulations reproduced both types of experimental data well: there was very good agreement for the SAXS profile over the entire *q* (momentum transfer) range from 0 to 0.5 Å^−1^; likewise the calculated secondary chemical shifts were close to the experimental values, with root-mean-square error (RMSE) at 0.43 ppm. The experimental SAXS profile is also reproduced well by the generated conformations, validating the latter’s sampling of the overall shape and size of ChiZ, though minor deviations are noticeable around *q* = 0 and around 0.3 Å^−1^. For secondary chemical shifts, the RMSE increases to 0.57 ppm for the generated conformations, likely reflecting violation of bond lengths and bond angles and over-smoothing of local backbone contours. For several residues near the two chain termini, the secondary chemical shifts are over-predicted. Still, the RMSE of 0.57 ppm for the autoencoder-generated conformations are lower than those (0.63 to 0.84 ppm) for conformations from MD simulations using several force fields^14^.

**Figure 7.**
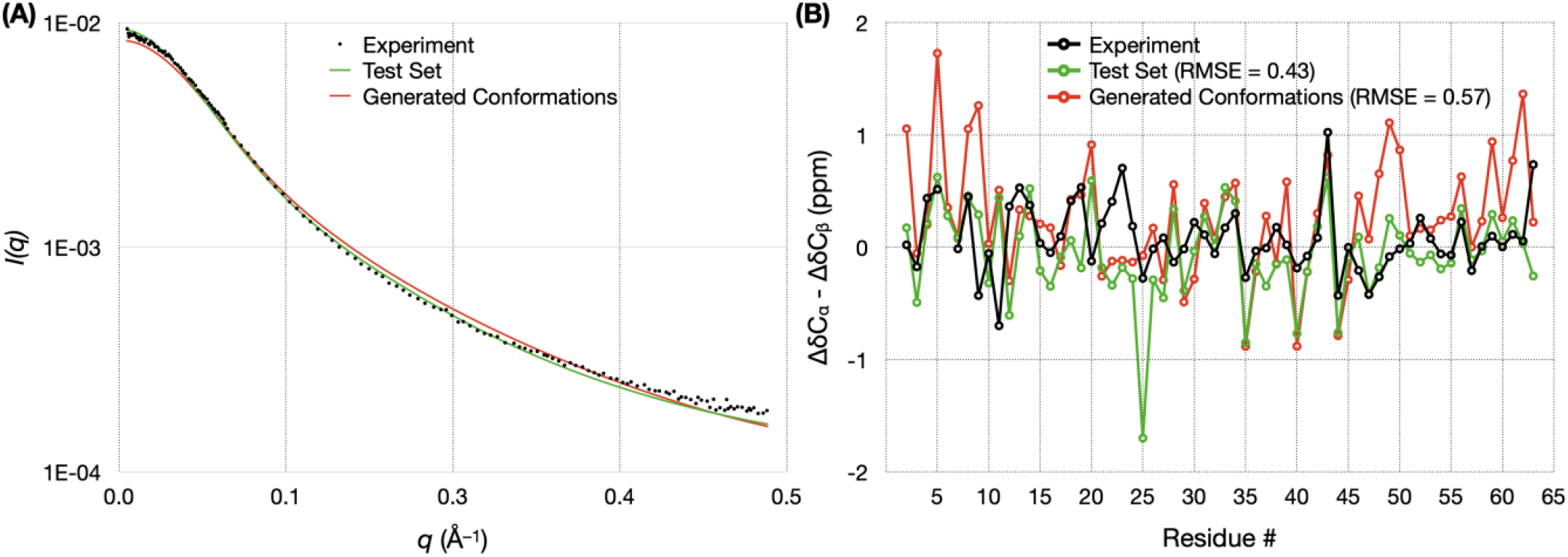
Validation of autoencoder-generated conformations for ChiZ by experimental SAXS and chemical shift data. (A) Comparison of experimental and calculated SAXS profiles. (B) Comparison of experimental and calculated secondary chemical shifts. The experimental data and the MD simulations are reported previously^14^. Calculations were done on either the test set comprising 12,180 conformations sampled from 12 MD runs, or on an autoencoder-generated set comprising the same number of conformations. The autoencoder was trained on a combined training set comprising 52,200 conformations sampled from the 12 MD runs.

## Discussion

We have developed generative autoencoders to mine the broad conformational space of IDPs. These autoencoders can not only represent IDP conformations in the latent space with high fidelity to allow for accurate reconstruction, but also generate new conformations to fill up the conformational space. The generated ensemble contains close matches for all the conformations sampled in long MD simulations, but with negligible computational time. For example, sampling 100,000 conformations (at 20-ps intervals) from MD simulations of Aβ40, even with GPU acceleration^21^, takes 80 days, whereas our autoencoder generates the same number of conformations in 12 seconds. In the case of ChiZ, the autoencoder-generated conformations even yielded better predictions for SAXS profile and chemical shifts than MD simulations with several force fields.

Our generative autoencoders have the flavor or variational autoencoders but are more intuitive. Rather than optimizing Gaussians in the latent space during the training process as in variational autoencoders, we only optimize reconstruction and then use the latent vectors of the training set to calculate the mean vector and covariance matrix, which are directly used to define a multivariate Gaussian for generating new conformations. At present the generated conformations have two shortcomings. First, they violate bond lengths and bond angles. Noé et al.^7^ appear to have found a working solution to this problem in building Boltzmann generators for structured proteins. It will be interesting to see whether that solution can be adapted for IDPs. Second, the backbone traces of the generated conformations are overly smooth locally. Over-smoothing is a common problem in many neural network models; ideas introduced in other fields (e.g., speech synthesis^22^) may provide inspiration.

The generative autoencoders designed here are for mining the conformational space of IDPs in isolation. The power of this approach demonstrated here suggests that it can be extended to study IDPs in more complex functional states, such as when bound to or associated with an interaction partner (a target protein or a membrane), or in aggregation. For example, ChiZ associated with acidic membranes has been studied by long MD simulations^15^; generative autoencoders may also be able to mine the conformational space of membrane-associated IDPs. IDPs are prone to phase separation^23^, resulting in a highly concentrated phase surrounded by a dilute phase. Microsecond-long MD simulations failed to sample the equilibration between the two phases^24^. AI-based models such as generative autoencoders may open the door to solving this and other challenging conformational mining problems for IDPs.

## Computational Methods

### Autoencoder design

We built and trained the autoencoders using the Keras package (https://keras.io/) with TensorFlow backend (https://www.tensorflow.org/) in Python 3.6^25^. The autoencoders consisted of an encoder and a decoder. Both the encoder and decoder had a dense neural network architecture, with two hidden layers of 300 and 50 neurons, respectively. The input, hidden, and output layers of the encoder and decoder were arranged as mirror images of each other (Fig. 1A). This arrangement was chosen based on its reduced training complexity as shown in previous reconstruction work on structured proteins^6^. All layers except for the final output layer had a rectified linear unit activation function; the final output layer had a sigmoidal activation function. The loss function was cross entropy, and the neural networks were trained by the Adam optimizer given its effectiveness in handling large datasets. For each autoencoder, training was done for 100 epochs using a batch size of 40.

The input to the encoder consisted of the Cartesian coordinates of an IDP. Only heavy atoms (all for the backbone and selected for side chains) were included; selected side-chain atom types were CB, CG, CD, OE1, and NE2. This selection contained all the heavy atoms of polyglutamine Q15, but truncated some of the side chains in Aβ40 and ChiZ. Q15, Aβ40, and ChiZ had *N* = 140, 230, and 385 heavy atoms, respectively, for a total of 3*N* input coordinates. The dimensions of the latent spaces for the three IDPs were 13, 30, and 40.

The latent space dimension and training size were tested based on reconstruction, which entailed encoding (i.e., representing the conformations as vectors in the latent space) and then decoding (i.e., constructing back full conformations from the latent vectors). Parameters of autoencoders trained on reconstruction were saved in decoder and encoder files, and the decoder was then used to generate new conformations.

### Molecular dynamics simulations

Two 1-μs trajectories (100,000 frames each, saved at 10-ps intervals) for Q15, taken from Hicks and Zhou^20^, were run in GROMACS with the Amber ff03ws force field^26^ for protein and TIP4P/2005 for water^27^. Twelve 3-μs trajectories (150,000 frames each, saved at 20-ps intervals) for ChiZ, taken from Hicks et al.^14^, were run on GPUs using pmemd.cuda^21^ in AMBER18^28^ with the ff14SB force field^29^ for protein and TIP4P-D^30^ for water. The protocol for ChiZ was used to run four replicate simulations of Aβ40 (3.5 μs each; 175,000 frames saved at 20-ps intervals).

### Data preprocessing

MD trajectories in GROMACS and AMBER trajectory formats were first converted to conformations in PDB format (with solvent stripped). An initial portion of each trajectory (5000, 35000, and 5000 frames for Q15, Aβ40, and ChiZ, respectively) were removed. The remaining trajectory was split into two parts, the first (e.g., 10%) as the training dataset and the second as the test dataset.

The Biobox library in Python (https://github.com/degiacom/biobox)^6^ was used to preprocess the coordinates in each dataset. All the frames were aligned to the first one according to RMSD, and shifted to have all coordinates positive. Coordinates were then scaled between 0 and 1 (via dividing by the maximum coordinate value) for using as input to the encoder. The output coordinates of the decoder were scaled back to real coordinates using the same scaling factor.

### RMSD calculation

We used a code of Ho (https://boscoh.com/protein/rmsd-root-mean-square-deviation.html) to calculate RMSDs of output conformations. A custom Python code (https://github.com/aaayushg/generative_IDPs/tree/main/RMSD) was written to find the lowest RMSD between a given test conformation against a set of generated conformations, and calculate the average of these best-match RMSDs for the test set (100-fold diluted).

### Generating new conformations

The mean vector and covariance matrix of the training dataset in the latent space were calculated to define a multivariate Gaussian distribution (Fig. 1B), from which vectors were sampled and fed to the decoder to generate new conformations (Fig. 1C). The multivariate Gaussian sampling was implemented using the NumPy library (https://numpy.org/) in Python. Histograms were calculated in two-dimensional subspaces of the latent space, for comparison among the training, test, and multivariate Gaussian datasets (https://github.com/aaayushg/generative_IDPs/tree/main/Plot_histogram).

### Calculation of SAXS profile and chemical shifts for ChiZ

Side-chain heavy atoms truncated for implementing autoencoders were added back using the FoldX program^31^. The SAXS profile for each conformation was then calculated using FoXS^32^ and scaled to optimize agreement with the experimental profile^14^. Chemical shifts were calculated using SHIFTX2^33^ (www.shiftx2.ca). Chemical shifts for random-coil conformations calculated using POTENCI^34^ were subtracted to obtain secondary Cα and Cβ chemical shifts. SAXS profiles and chemical shifts were averaged over all the conformations in the diluted test set (12180 frames from 12 trajectories) or a generated set (of the same size).

## Data and code availability

The implementation codes, saved autoencoder models, tutorials, and example data are available on GitHub: https://github.com/aaayushg/generative_IDPs.

## Acknowledgment

This work was supported by National Institutes of Health Grant GM118091.

