## Supplementary Table and Figures for "Artificial Intelligence Guided Conformational Mining of Intrinsically Disordered Proteins"

*Supporting Information*

**Table S1. Diversity of test sets and similarity between training and test sets**

| | Pairwise RMSD (Å)[a] (dil test × dil test) | Best Match RMSD (Å)[b] (dil test × dil test) | Best Match RMSD (Å)[c] (dil test × train) |
|---|---|---|---|
| Q15 (10% run1) | 6.98 | 3.71 | 3.96 |
| Aβ40 (20% run1) | 11.61 | 3.83 | 6.76 |
| ChiZ (30% run1) | 18.21 | 4.83 | 10.17 |
| ChiZ (combined)[d] | 19.23 | 8.62 | 8.47 |

[a]Average RMSD when each conformation is compared with all other conformations in the diluted test set.

[b]Average best-match RMSD of the diluted test set against other members of the same set.

[c]Average best-match RMSD of the diluted test set against the training set.

[d]Combined training or test set, each from combining the corresponding data from all of the 12 MD runs. The combined training set is diluted 10-fold before use, whereas the combined test set is diluted 1000-fold.

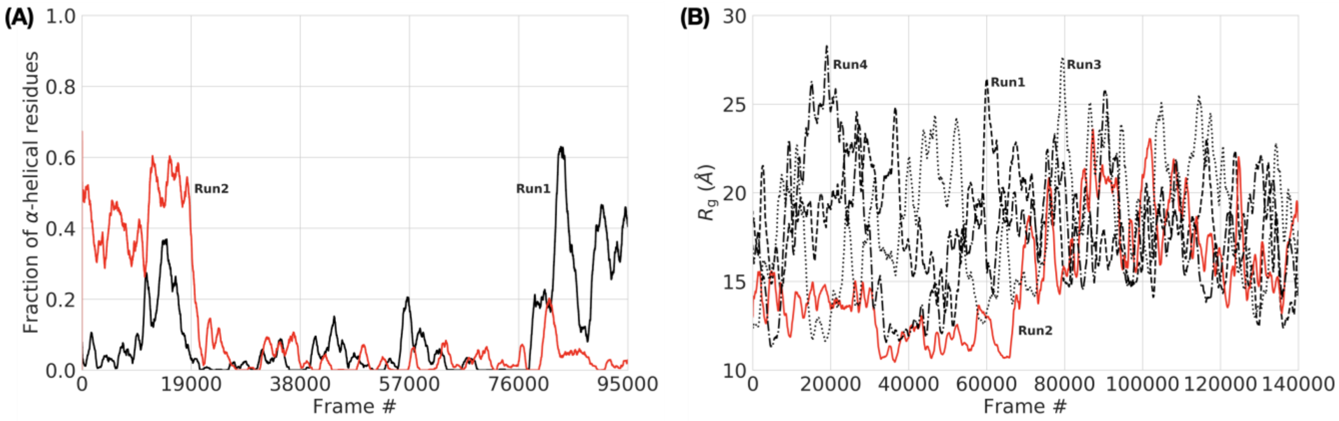


**Figure S1. Conformational properties of Q15 and Aβ40 in MD simulations.** (A) Fraction of α-helical residues in two replicate MD runs of Q15. Run1 started from a random-coil conformation whereas run2 started from an all α-helical conformation. These simulations were reported previously[1], where conformational sampling was enhanced by the replica exchange with solute tempering (REST2) method[2,3]. (B) Radius of gyration of Aβ40 in four replicate runs. All the four runs started from disordered conformations, but run2 happened to stay relatively compact for the first half of the simulation. The curves were smoothed by running average with a window of 1000 frames (sampled at 10-ps intervals for Q15 and 20-ps intervals for Aβ40).

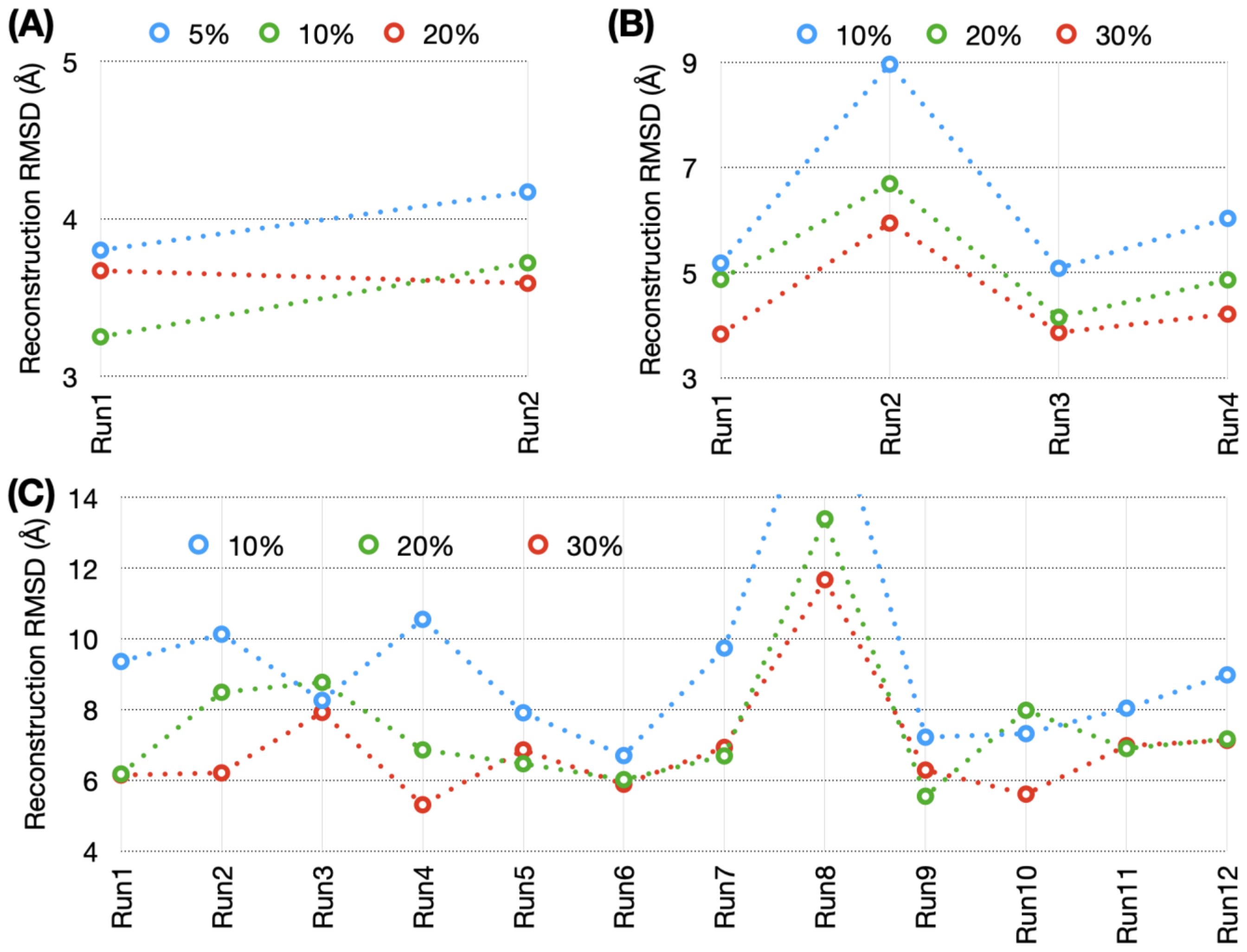


**Figure S2. Reconstruction results when the dimension of the latent space is increased to 200.** Average reconstruction RMSDs at different sizes of the training sets sampled from replicate MD runs. (A) Q15 at 5%, 10%, and 20% training sizes from two runs. (B) Aβ40 at 10%, 20%, and 30% training sizes from four runs. (C) ChiZ at 10%, 20%, and 30% training sizes from 12 runs.

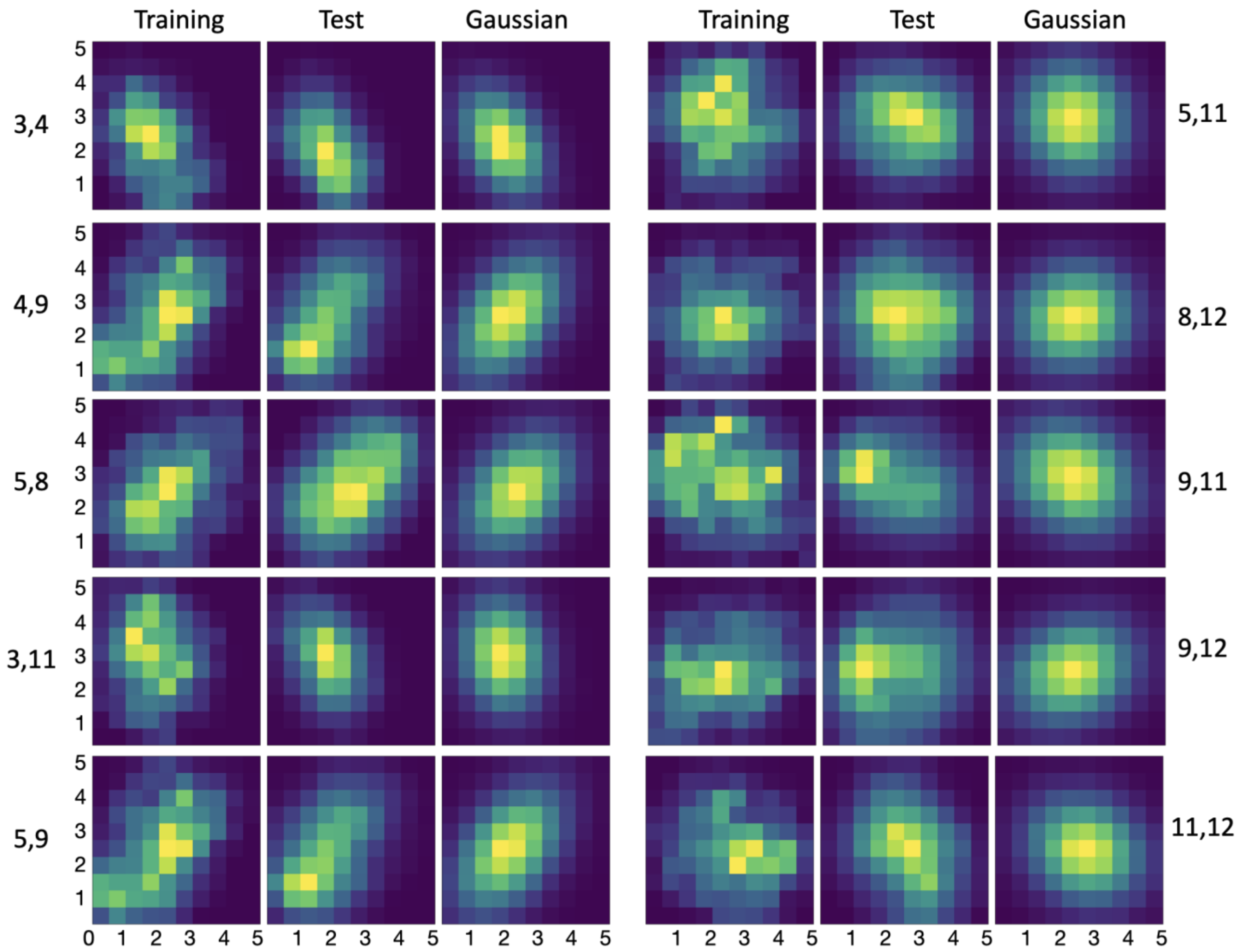


**Figure S3. Histograms of Q15 in the latent space, calculated from training data, test data, and multivariate Gaussian.** Histograms for pairs of encoder nonzero outputs from run1 are shown as heat maps, with yellow representing pixels with the highest counts and dark blue representing pixels with 0 count.

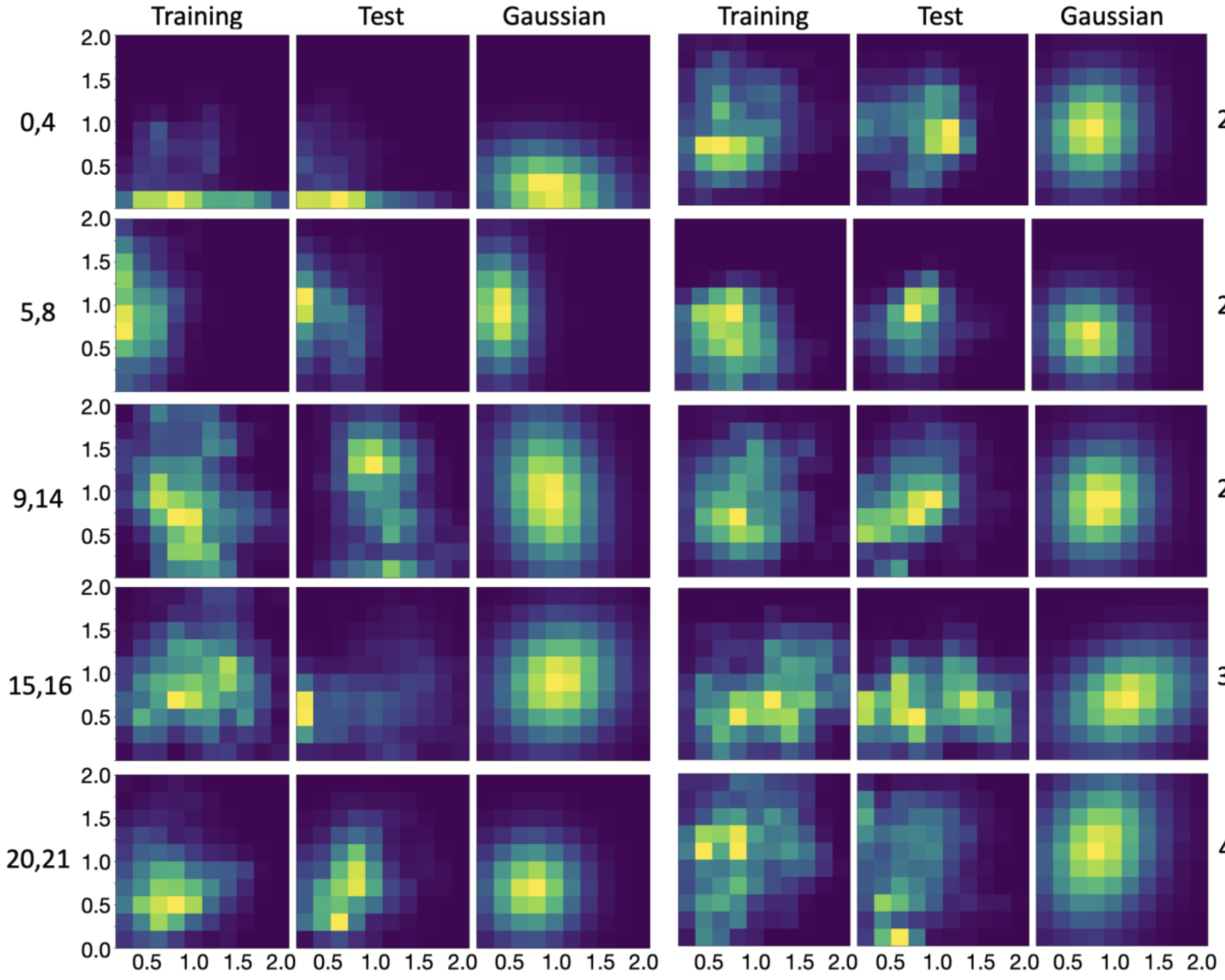


**Figure S4. Histograms of ChiZ in the latent space, calculated from training data, test data, and multivariate Gaussian.** Histograms for pairs of encoder nonzero outputs from run1 are shown as heat maps, with yellow representing pixels with the highest counts and dark blue representing pixels with 0 count.

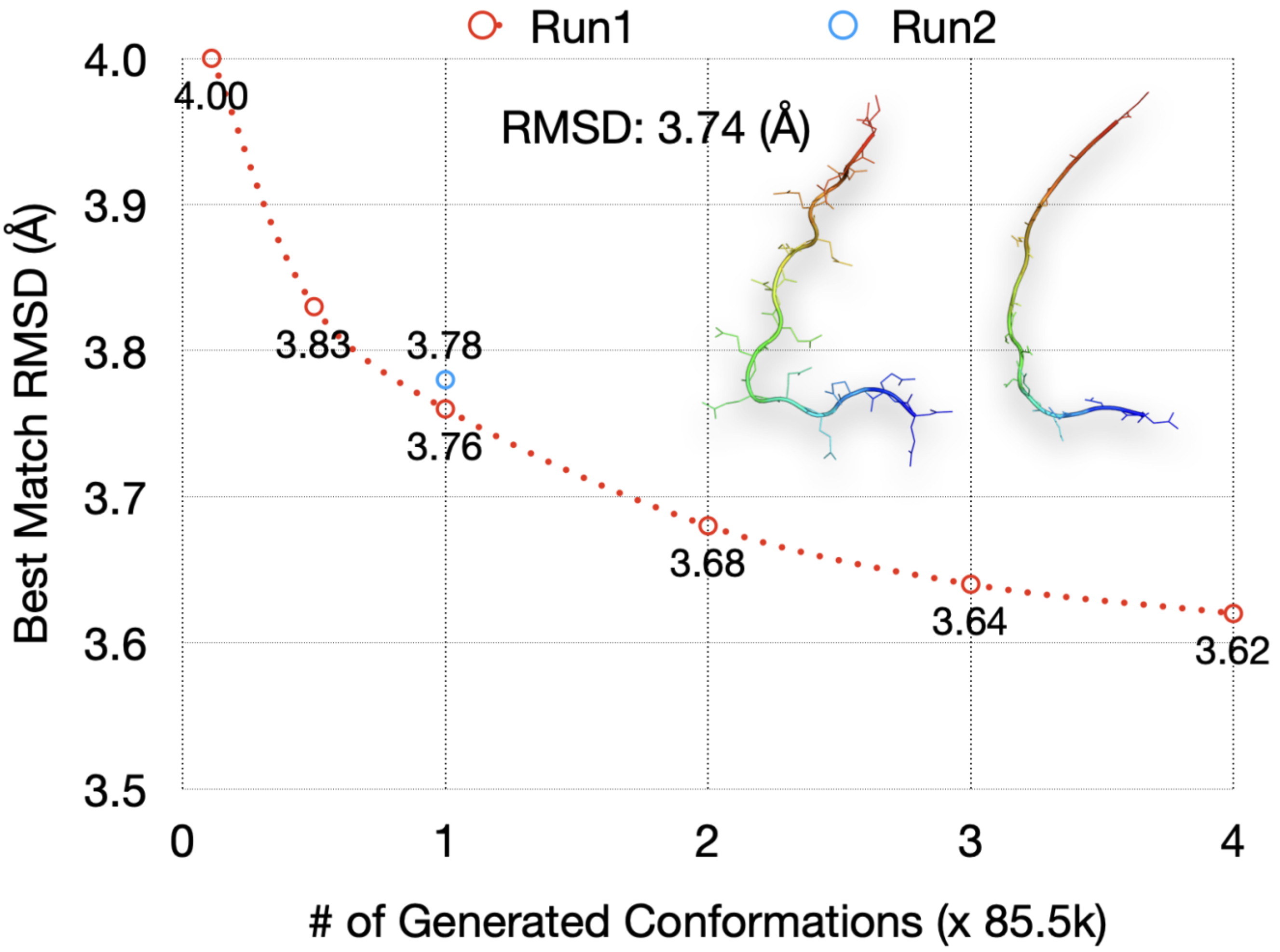


**Figure S5. Results for autoencoder-generated conformations of Q15, with a 200-dimensioan latent space.** The average best-match RMSDs of the 100-fold diluted test set are calculated against generated sets at different sizes. The latter sizes are measured in multiples of the test size (= 85,500). Run1 results are shown at sizes of the generated set ranging from the training size (9,500 or 0.11×) to 4×. For run2, the result is shown at 1×.

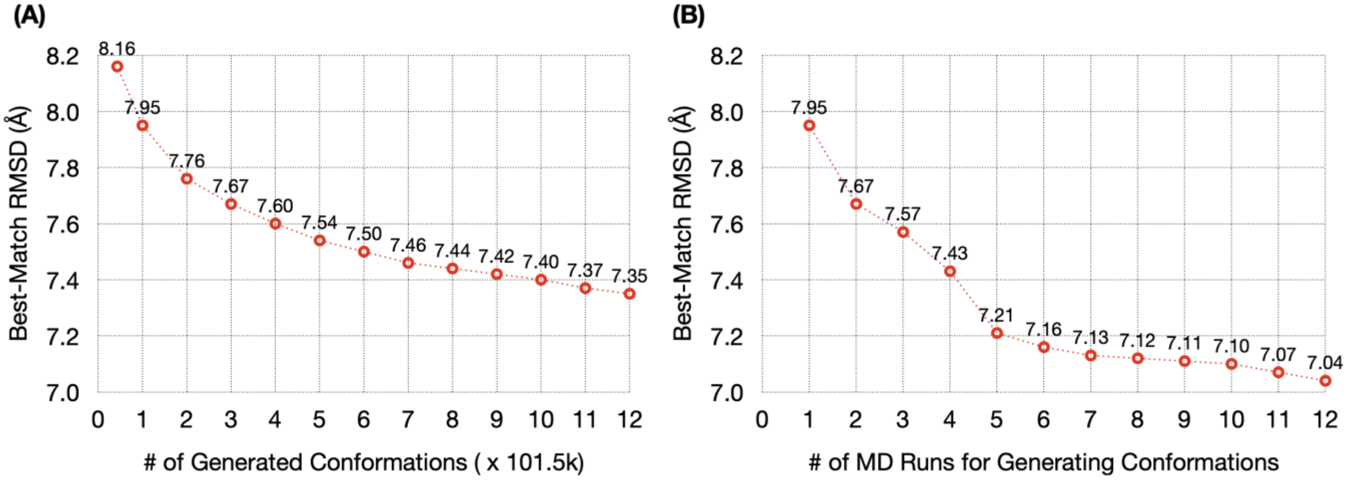


**Figure S6. Results for autoencoder-generated conformations of ChiZ.** The average best-match RMSDs of the 100-fold diluted test set of run1 are calculated against generated sets at different sizes. (A) Generated sets from run1, at sizes measured in multiples of the test size (= 101,500). (B) Generated sets pooled from run1 to run*i*, where *i* goes from 1 to 12. From each MD run, the generated set is at size 1×.
